# Comparative transcriptome analysis of Powdery mildew Resistance between two Melon (*Cucumis melo* L) with Different Thickness Peel

**DOI:** 10.1101/541391

**Authors:** Cheng Hong, Kong Wei-ping, Lü Jun-Feng

## Abstract

Melon (*Cucumis melo* L.) is wildly planted in the world and China is a major producer of muskmelon. Powdery mildew is one of the most common fungal diseases in the world and this disease frequently affects melon (*Cucumis melo* L.) and due to the reduction of melon yield. In this study, one material GanTianmi with thin peel and another material XueLianHua with thick peel were selected. After inoculating the powdery mildew, both materials were used to do the RNA-Seq. In total two RNA-seq libraries were constructed and sequenced separately. The reads per kilobase per Million mapped reads (RPKM) values of all the genes in the two materials were calculated and there were 13828 genes were expressed in the material G and 13944 genes were expressed in the material S (RPKM>1). The differentially expression gene (DEG) analysis result suggested that total 769 the DEGs between the two materials were identified. All the DEGs were annotated with several database and the transcript factors (TFs) that related to disease resistance such as *MYB*, *ERF* and *WRKY* among the DEGs were also identified. This research could not only provide the information about understanding the mechanism of powdery mildew infection but also help researchers breed the varieties with powdery mildew resistance.

## Introduction

Melon (*Cucumis melo* L.) is one of the most valuable cash crops that grown all over the word and hina also is a major producer of muskmelon (Liu, 2003; Chen et al, 2015). However, powdery mildew often infects muskmelon, thereby resulting in significant losses in its production (Cheng et al., 2006). In previous studies, there were lots of research that identified candidate genes/function genes about the powdery mildew resistance (Fukino et al, 2008; Chen et al, 2012, 2013 and 2015; Natarajan et al, 2016; Wang et al, 2016)

Some function genes were identified through forward genetics such as genetic map construction and quantitative trait locus (QTL) identification. The genes that located on the confidence intervals of the QTLs for powdery mildew resistance were considered as candidate genes/function genes (Fukino et al, 2008; Natarajan et al, 2016; Wang et al, 2016). There were also some that identified candidate genes/function genes for powdery mildew resistance through reverse genetics such as using mutant (Chen et al, 2012, 2013 and 2015). From these researches, there were also some that suggested the melon that with thin peel often manifested as disease resistance while the melon that with thick peel often manifested as disease susceptible.

In this study, one material GanTianmi with thin peel and another material XueLianHua with thick peel were selected. After inoculating the powdery mildew, both materials were used to do the RNA-Seq. In total, two RNA-seq libraries were constructed and sequenced separately; generating 47,462,015 mean clean reads with mean GC content of 49.32 and mean Q30 of 85.89%. Among all the genes in the genome of melon, 13828 genes were expressed in the material G and 13944 genes were expressed in the material S (RPKM>1). Through multiple comparisons between libraries, 769 DEGs were identified, of which the 726 were up-regulated DEGs and 43 were down-regulated ones. Based on the annotation information in gene ontology (GO), kyoto encyclopedia of genes and genomes (KEGG) and eukaryotic orthologous groups (KOG) database, most of the DEGs were related to “nucleus”, “chloroplast” and “cytoplasm” in GO database; related to “Metabolic pathways”, “Biosynthesis of secondary metabolites” and “Ribosome” in KEGG database and related to “Signal transduction mechanisms” “Posttranslational modification, protein turnover, chaperones” and “Lipid transport and metabolism” in KOG database. There were six genes that have annotation information of “Plant-pathogen interaction “. There were also 46 genes were TFs, among them, four were the *MYB* (Li et al, 2013; Wang et al, 2013; Yang et al, 2019), six were the *ERF* (Xiang et al, 2018; Debbarma et al, 2019) and two were the *WRKY* (Line et al, 2011; Jiao et al, 2018; Yang et al, 2018);. This TFs were all related to disease resistance (Ling et al, 2010; Li et al, 2013; Wang et al, 2013; Jiao et al, 2018; Xiang et al, 2018; Yang et al, 2018; Debbarma et al, 2019; Yang et al, 2019). All this research could provide information for breeding muskmelon cultivars that were resistant to powdery mildew.

## Materials and methods

### Plant materials

There were two materials were used in this research. One material was named GanTianmi (G). This material was manifested as disease resistance with thin peel. The other material was named XueLianHua (H) was manifested as disease susceptible with thick peel.

### RNA sample collecting and extraction

The cultivars were sown in a plastic nutrition pot after accelerated germination, and then place in a box under artificial climate, with a night temperature of 18 °C and a day tempera- ture of 25 °C. When the seedlings reached the Three-leaf stage, the powdery mildew germ Race1 was collected using a single lorica (Podosphaera xanthii), which is widely distributed in the Gansu Province. The seedlings were then infected with the plant pathogen in a bioclean room by using the leaf inoculation method. Three days after inoculation, spots of powdery mildew on the blades of the seedlings were collected and used for RNA extraction. Total RNA was extracted by using the TRIzol reagent (Beijing Tiangen), following the manufacturer’s instructions.

### RNA library preparation, sequencing, and quality control

The RNA samples were used to generate sequencing libraries according to the manufacturer of Illumina TruSeq™ RNA Sample Preparation Kits (Illumina, San Diego, CA, USA) Transcriptome sequencing was carried out on an Illumina HiSeq 2500 platform that produced 150 base pair (bp) paired-end (PE) raw reads (Personalbio, Shanghai, China). Raw data (raw reads) in FASTQ format were processed used the trimmomatic software (Bolger *et al.*, 2014). In this step, clean data were obtained as follows: the first, removing reads with ≥10% unidentified nucleotides (N); the second, Removing reads with > 50% bases having phred quality < 5; the third, Removing reads with > 10 nt aligned to the adapter, allowing ≤10% mismatches; the fourth, removing putative PCR duplicates ≤ generated by PCR amplification in the library construction process. After trimmomatic, the quality of the clean data was checked by the software FASTQC (http://www.bioinformatics.babraham.ac.uk/projects/fastqc/).

### Read mapping and expression level analysis

The clean sequence tags were mapped to the *Cucumis melo* (http://cucurbitgenomics.org/organism/3). An index of the reference genome was built using Bowtie v2.0.6 (Langmead et al, 2012) and PE clean reads were aligned to the reference genome using TopHat v2.0.9 (Kim et al, 2013). The reads per RPKM values of genes were calculated using the software of cufflinks program (Trapnell et al., 2010). After got the RPKM of the genes, the expression heatmap of the genes were drawn with the R package Pheatmap with the value of log2 (RPKM+ 1).

### Gene expression pattern analysis

The gene expression level was estimated by counting the number of reads that could be map to genes or exons in the reference genome. To make gene expression data coincident and could be comparable across different genes and experiments, the RPKM was used. The gene expression levels were analyzed with software HTSeq (https://htseq.readthedocs.io/en/release_0.10.0/), model “union”. Differential expression analysis of two genes was analyzed with the software DESeq in the R package. This software could provide statistical routines to determine differential expression in digital gene expression data using a model based on the negative binomial distribution The genes that with q_value less than 0.05 and the log2 (Fold_change) (log2 (FC)) more than one or less than −1 could be considered as DEGs. The genes that with log2 (FC) more than one were up-regulated genes while ones with log2 (FC) less than negative one were down-regulated genes.

### Different expressed genes annotation and transcription factor analysis

The GO annotation information of the candidate genes was got from the GO databases (http://archive.geneontology.org/latest-lite/) and (ftp://ftp.ncbi.nlm.nih.gov/gene/DATA/). The software BLAST was used to compare the sequence of the genes and the sequence in the GO database. The KEGG annotation information was got through the software KOBAS 3.0 (Xie et al, 2010). The KOG annotation information of candidate genes was got from the KOG database (ftp://ftp.ncbi.nih.gov/pub/COG/KOG). The rules of comparison were same as it in GO analysis. All the transcripts of melon were downloaded in the database of Plant TFDB (http://plantTFDB.cbi.pku.edu.cn/) and then the DEGs were compared with these TFs with the NCBI Blast. The sequence results with *E_value* less then 1e-10 could be considered as TFs.

## Results

### Transcriptome sequencing and alignment to the reference genome

With the purpose of systematic identification of the candidate genes that related to the difference in disease resistance between the materials G and H, RNA-seq libraries of fiber samples were constructed and analyzed. For the materials G, there were 56,645,955raw reads, total 16,540,359,199bp, 49.24% GC content and the Q30 was 86.02%; after trimmomatic, there were 48,482,165 clean reads, total 137692342bp. For the materials H, there were 54343040 raw reads, total 15,812,206,404bp, 49.4% GC content and the Q30 was 49.40%; after trimmomatic, there were 46441864 clean reads, total 13,113,637,322bp (Table S1).

In the alignment analysis, the proportion of clean data was mapped to the genome of *Cucumis melo.* For the materials G, there were 87,830,047 (90.58%) reads that could be uniquely mapped to the genome of *Cucumis melo* and 87001260 of them could be mapped to genes (99.60% in exon). For the materials H, there were 82,313,164 (88.62%) reads that could be uniquely mapped to the genome of *Cucu*mis melo and 80435970 of them could be mapped to genes (99.59% in exon) (Table S2).

### The expression level of the genes

From the RNA sequencing, all the expression levels of the genes in the *Cucumis melo* genome were calculated. For the materials G, total 7222 genes were not expressed as the RPKM of these genes were zero; total 6377 genes were low expressed as the RPKM of these genes were between zero and one; total 8944 genes were medium expressed as the RPKM of these genes were between one and ten; total 4884 genes were high expressed as the RPKM of these genes were more than 10; For the materials H, total 7275 genes were not expressed; total 6237 genes were low expressed; total 9103 genes were medium expressed a; total 4841 genes were high expressed (Table S3, Table 1, Figure 1).

**Table 1.** The Basic statistics of RPKM_value of the genes in two materials

| RPKM Value | G | H |
| --- | --- | --- |
| RPKM=0 | 7222 | 7245 |
| 0<RPKM<1 | 6377 | 6237 |
| 1<RPKM<10 | 8944 | 9103 |
| RPKM>10 | 1884 | 4842 |

**Table 2.**
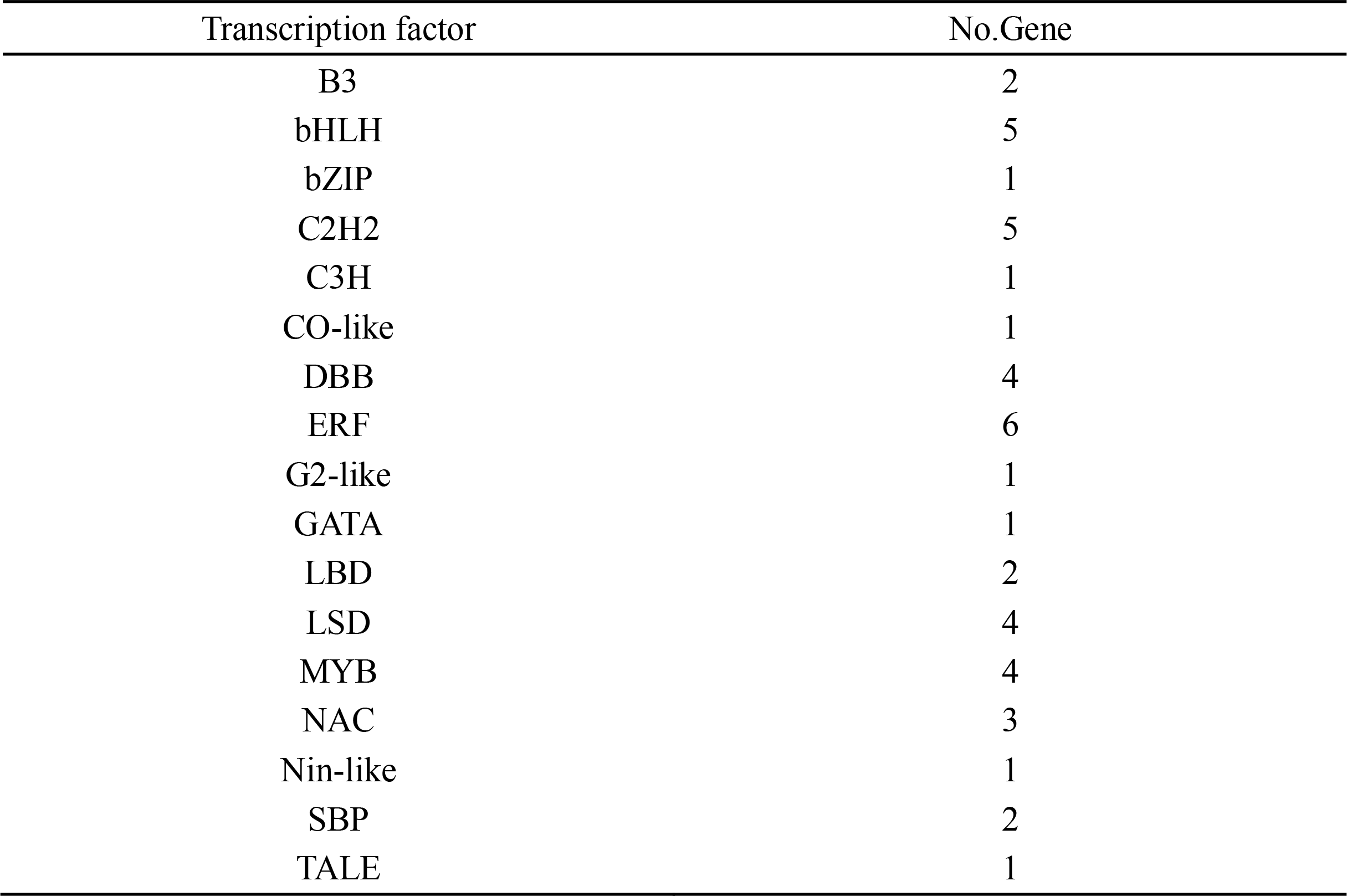
The number of TFs in the DEGs

| Transcription factor | No.Gene |
| --- | --- |
| B3 | 2 |
| bHLH | 5 |
| bZIP | 1 |
| C2H2 | 5 |
| C3H | 1 |
| CO-like | 1 |
| DBB | 4 |
| ERF | 6 |
| G2-like | 1 |
| GATA | 1 |
| LBD | 2 |
| LSD | 4 |
| MYB | 4 |
| NAC | 3 |
| Nin-like | 1 |
| SBP | 2 |
| TALE | 1 |

**Figure 1.**
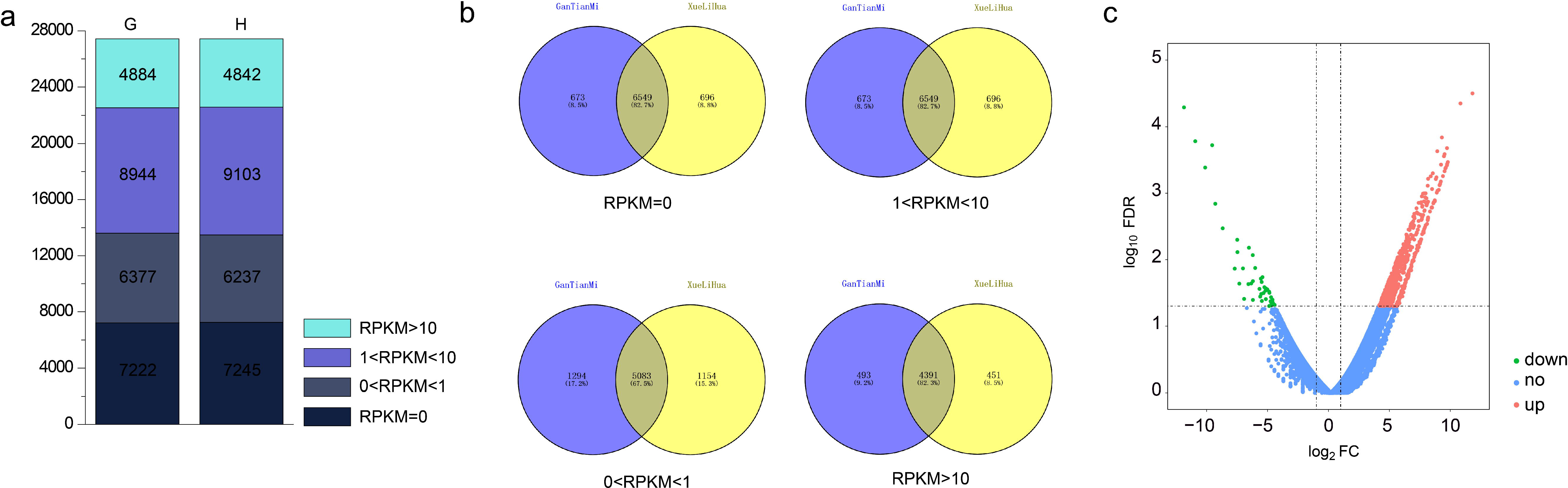
a. The number of genes that with RPKM intervals of RPKM=0, 0<RPKM<1, 1<RPKM<10, RPKM>10 b. The intersection and union of the genes that with RPKM intervals of RPKM=0, 0<RPKM<1, 1<RPKM<10, RPKM>10 between the material G and H c. The log2FoldChange value and FDR value of the positive regulation DEGs and negative regulation DEGs

For the no expressed genes there were 6549 genes (82.7%) that could be detected in both materials, 673 could be detected in only G and 696 could be detected in only H; for the weak expression genes, there were 5083 (67.5%) could be detected in both materials, 1294 (17.2%) could be detected in only G and 1154 (15.3) could be detected in only H; for the medium expressed genes, there were 7932 could be detected in both materials, 1012 (10.0%) could be detected in only G and 1171 (11.6%) could be detected in only H; for the high expressed genes, there were 4391 (82.3%) could be detect in both materials, 493 could be detected in only G and 451 could be only detected in H (Table S3, Table 1, Figure 1).

### The different expression genes

To find out the different expression genes between the two materials G and H, the software Deseq2 were used and the genes with the absolute value of log2FC more than one and Q_value less than 0.05 could be considered as different expression genes. In this research, there were 769 different expression genes. Among them, 726 were up-regulation genes and the other 43 were down-regulation genes (Table S4, Figure 2).

**Figure 2.**
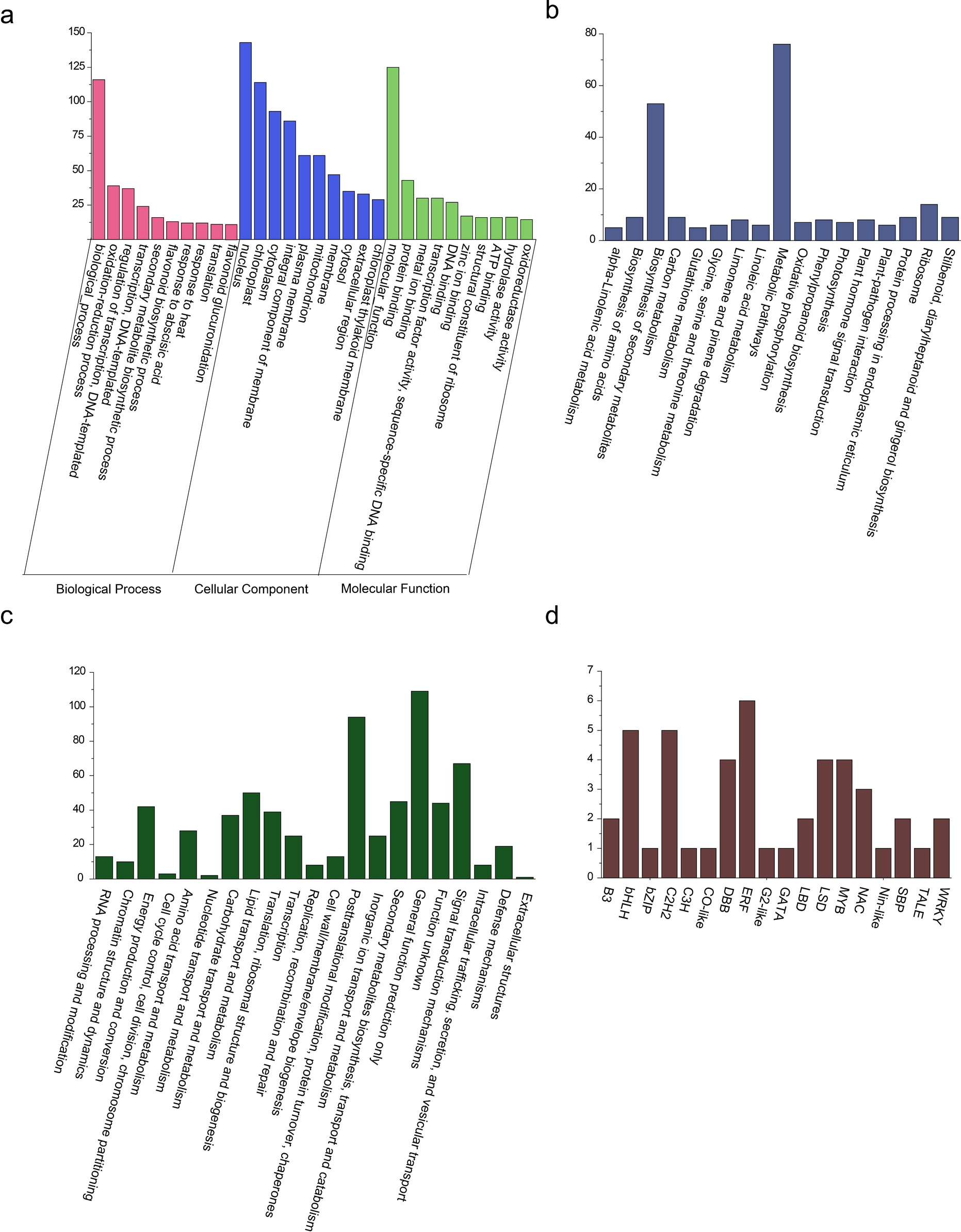
The annotation information of the DEGs between the materials G and H a. The GO annotation information of the DEGs between the materials G and H b. The KEGG annotation information of the DEGs between the materials G and H c. The KOG annotation information of the DEGs between the materials G and H d. The TF annotation information of the DEGs between the materials G and H

### The annotation with different expression gene

All the different expression genes were annotated with GO, KEEG and KOG. For the GO annotation, all the DEGs could be annotated into 968 GO terms as one DEG maybe annotated into more than one GO term. Among the 968 GO terms, 326 belonged to the category of molecular function; 146 belonged to the category of cellular component and 496 belonged to the category of biological process. The GO term nucleus harbored the most number of DEGs (143), and followed with the GO term molecular function, biological process and cytoplasm, harbored 125 116 and 114 DEGs respectively (Table S5, Figure 2). For the KEGG analysis, all the DEGs were annotated into 76 KEGG pathways. Among them, “Metabolic pathways”, “Biosynthesis of secondary metabolite” and “Ribosome” harbored the most DEGs (76, 53 and 14) respectively (Table S6, Figure 2); For the KOG analysis, all the DEGs were located into 21 KOG baskets. Among them, the basket “Signal transduction mechanisms”, “Lipid transport and metabolism” and “Energy production and conversion “harbored the most DEGs (67, 50 and 42) respectively. There were also 153 DEGs that with unclear KOG annotation information as 109 of them located on the KOG basket of “General function prediction only” and the other 44 located on the KOG basket of “Function unknown” (Table S7, Figure 2).

### Transcription factor analysis

ALL The DEGs were compared with the TFs in the database of (http://planttfdb.cbi.pku.edu.cn/). Totally, 46 DEGs could be considered as TFs. Among them, there were six DEGs that were *ERF* TF, four were *MYB* TF and two were *WARK* TF. The detail information about TF analysis was in the Figure 2 and Table S8. The TFs *ERF*, *MYB*, and *WARK* were related to disease resistance (Table S8, Figure 2).

## Discussion

From the annotation information of the DEGs, there were six genes that annotated with the KEGG pathway “Plant-pathogen interaction”. The six DEGs were *MELO3C016746, MELO3C015295, MELO3C021706, MELO3C018539, MELO3C020725* and *MELO3C000209.* Among them, the DEG *MELO3C018539* have been published that it played the role of against *cucurbit chlorotic yellows virus* in melon (Takeshita et al, 2013), but the other five DEGs have not been published in melon. In the *Arabidopsis thaliana* annotation database, the homologous gene of *MELO3C016746* in *Arabidopsis thaliana* (*AT1G29150*) was specifically interacted with FUS6/COP11 via the C-terminal domain of *FUS6/COP11* and associates with an ATPase subunit of the 19S proteasome regulatory complex, AtS6A. The mRNA is cell-to-cell mobile (Cho et al, 2006 and 2015;); the homologous gene of *MELO3C015295* in *Arabidopsis thaliana* (*AT5G56010*) was a member of heat shock protein 90 (HSP90) gene family and expressed in all tissues and abundant in root apical meristem, pollen and tapetum (Cha et al, 2013; Bologna et al, 2018); the homologous gene of *MELO3C021706* in *Arabidopsis thaliana* (*AT3G18990*) was related to regulation of flower development and vernalization response (Kinjo et al, 2012; King et al, 2013); the homologous gene of *MELO3C020725* in *Arabidopsis thaliana* (*AT2G37650*) was belong to *GRAS* family transcription factor and it related to biological processes in regulating root and shoot development, plant growth, development and stress responses (Liu et al, 2018; Zhang et al, 2018); the homologous gene of *MELO3C000209* in *Arabidopsis thaliana* (*AT5G56030*) was also a member of heat shock protein 90 (HSP90) gene family and the function was similar with the gene *MELO3C015295* (Cha et al, 2013; Bologna et al, 2018).

There were also some DEGs that were TFs that related to disease resistance like *MYB* (Katlyar et al, 2012; Li et al, 2013; Wang et al, 2013; Yang et al, 2019), *ERF* (Xiang et al, 2018; Debbarma et al, 2019) and *WRKY* (Line et al, 2011; Jiao et al, 2018; Yang et al, 2018). In plants, MYB TFs were related to plant development, secondary metabolism, hormone signal transduction, disease resistance and abiotic stress tolerance (Katlyar et al, 2012); the ERF TFs plays an important role in plant development processes and stress responses such as salt, drought, heat, and cold with well-conserved DNA-binding domain (Xiang et al, 2018; Debbarma et al, 2019); The *WRKY* TFs were widely implicated in defense responses and various other physiological processes (Yang et al, 2018); it could evolve diverse functions with different biochemical properties and implicated to modulate seed development, flowering, fruit ripening, senescence and various metabolic processes (Jiao et al, 2018).

In our research, these DEGs were related to disease resistance and could be considered as candidate genes. But whether these DEGs could make some contributions to increase the disease resistance of melon, it still need further studies like transgene.

## Acknowledgements

The National Natural Science Foundation of China(31471896, 31360434) Technology Support Project of the GAAS (2017GAAS48) and The Major Project of Gansu Science & Technology (17ZD2NA015-04) supported this study.

## Supporting information

Table S1. The detail information of RNA-Seq quality evaluate

Table S2. The detail information of comparison efficiency evaluate

Table S3. The RPKM value of all the genes in the material G and H

Table S4. The log2FoldChange value and FDR value of the DEGs between the materials G and H

Table S5. The detail GO annotation information of DESs between the materials G and H

Table S6. The detail KEGG annotation information of DESs between the materials G and H

Table S7. The detail KOG annotation information of DESs between the materials G and H

Table S8. The detail TF annotation information of DEGs between the materials G and H

